# An optimized method for the isolation and the molecular characterization of cerebral microvessels *in vivo*

**DOI:** 10.1101/335224

**Authors:** Yun-Kyoung Lee, Helen Smith, Hui Yuan, Akira Ito, Teresa Sanchez

**Author notes:** Correspondence: Teresa Sanchez, PhD, Weill Cornell Medicine, Department of Pathology and Laboratory Medicine, 1300 York Ave, Box 69, A607B, New York, NY, 10065, USA. Equal contributions.

## Abstract

The molecular characterization of cerebral microvessels in experimental disease models has been hindered by the lack of a standardized method to reproducibly isolate intact cerebral microvessels, with consistent cellular compositions, and without the use of enzymatic digestion, which causes undesirable molecular and metabolic changes. Herein, we describe an optimized method for microvessel isolation from mouse brain cortex, which yields microvessel fragments (diameter <50 μm, 89.3% 3-5 μm) with consistent populations of discrete blood-brain barrier components (endothelial cells, pericytes, and astrocyte end feet), retaining high RNA integrity and protein postranslational modifications (e.g. phosphorylation). We demonstrate that this method allows the quantification of changes in gene expression in a disease model (stroke) and the activation of signalling pathways in mice subjected to drug administration. We also describe the isolation of genomic DNA and bisulfite treatment for the assessment of DNA methylation, as well as the optimization of chromatin extraction and shearing from cortical microvessels. Therefore, this protocol will be of great use to improve the understanding of the molecular mechanisms governing cerebrovascular dysfunction, which may help the development of novel therapies for stroke and other neurodegenerative diseases.

## INTRODUCTION

The cerebrovascular endothelium is a highly specialized vascular bed, which, in coordination with pericytes, vascular smooth muscle cells and glial cells forms a physical, transport and metabolic barrier (the blood brain barrier), maintains an anticoagulant and anti-inflammatory state, and regulate microcirculatory flow in order to meet neuronal demands and maintain homeostasis. Increasing preclinical and clinical evidence indicate that these fundamental functions of the endothelium are severely compromised in cerebrovascular and neurodegenerative diseases and significantly contribute to the exacerbation of neuronal injury in these conditions^1-5^. In addition, the therapeutic potential of the cerebrovascular endothelium is also well established, since it is positioned right at the interface between blood and brain parenchyma. However, despite extensive research, our understanding of the molecular mechanisms underlying cerebral microvascular dysfunction is still very limited.

Working towards this goal, several studies have attempted to characterize the molecular signature of the cerebrovascular endothelium by transcriptome and proteome profiling of isolated cerebral endothelial cells or mural cells ^6-9^. These approaches have allowed the identification of molecules enriched in the cerebrovascular endothelium compared to the endothelium of other organs or to other components of the neurovascular unit. In order to investigate the activation of signaling pathways upon pharmacological treatments or upon ischemic or inflammatory challenge, numerous *in vitro* studies with cultured brain endothelial cells have been conducted ^2,10,11^. However, both the unique phenotype and molecular signature of the endothelium from specific organs are lost upon isolation of the cells from their tissue microenvironment, deprivation from blood flow and in *vitro* culture^12,13^. Such protocols also prohibit the accurate study of junctional protein expression, post-translational modifications and subcellular localization in health and disease states within a living organism. For all these reasons, in order to conduct the molecular characterization of cerebral microvessels in disease models or upon pharmacological treatments the ability to study the intact microvasculature – isolated from the brain parenchyma while retaining the integrity of intercellular connections between the endothelium and mural cells – is a vitally useful tool. Various methods have been used to isolate microvessel fragments preserving the interactions of the endothelium with mural cells. Many of these methods use enzymatic digestion to achieve purity and consistency between preparations^14 15^, causing undesirable molecular and metabolic changes, compromising RNA and protein integrity and impairing the ability to detect low abundance transcripts, changes in gene expression or the activation of signaling pathways. In order to overcome these challenges, we have established a method for mouse cortical microvessel isolation (Figure 1), by combining, optimizing and supplementing various previously published protocols ^16,17 11^. The protocol described in this article yields intact microvessel fragments (diameter <50 μm, 89.3% 3-5 μm) with consistent populations of discrete blood-brain barrier components (endothelial cells, pericytes and astrocyte end feet) without a requirement for enzymatic digestion (Figure 2), retaining high RNA integrity (Figure 3) and protein postranslational modifications. We subsequently demonstrate that, the described protocol can be applied to quantify changes in gene expression in a disease model (stroke)^18^ (Figure 4) and the activation of signalling pathways in mice subjected to drug administration (Figure 5). In addition, we describe other successful applications of this protocol, useful for further molecular characterization of cerebral microvessels, such as the determination of DNA methylation status by isolation of genomic DNA, bisulfite treatment and methylation specific PCR (Figure 6), as well as the extraction of chromatin and shearing to the appropriate size for chromatin immunoprecipitation (ChIP) assays (Figure 7). Finally, we have recently published a study using the described protocol, in which quantitative changes in subcellular localization of tight junction proteins in cerebral microvessels from endothelial-specific genetically engineered mice and correlation with *in vivo* blood brain barrier and neuronal function are demonstrated ^19^.

**Figure 1.**
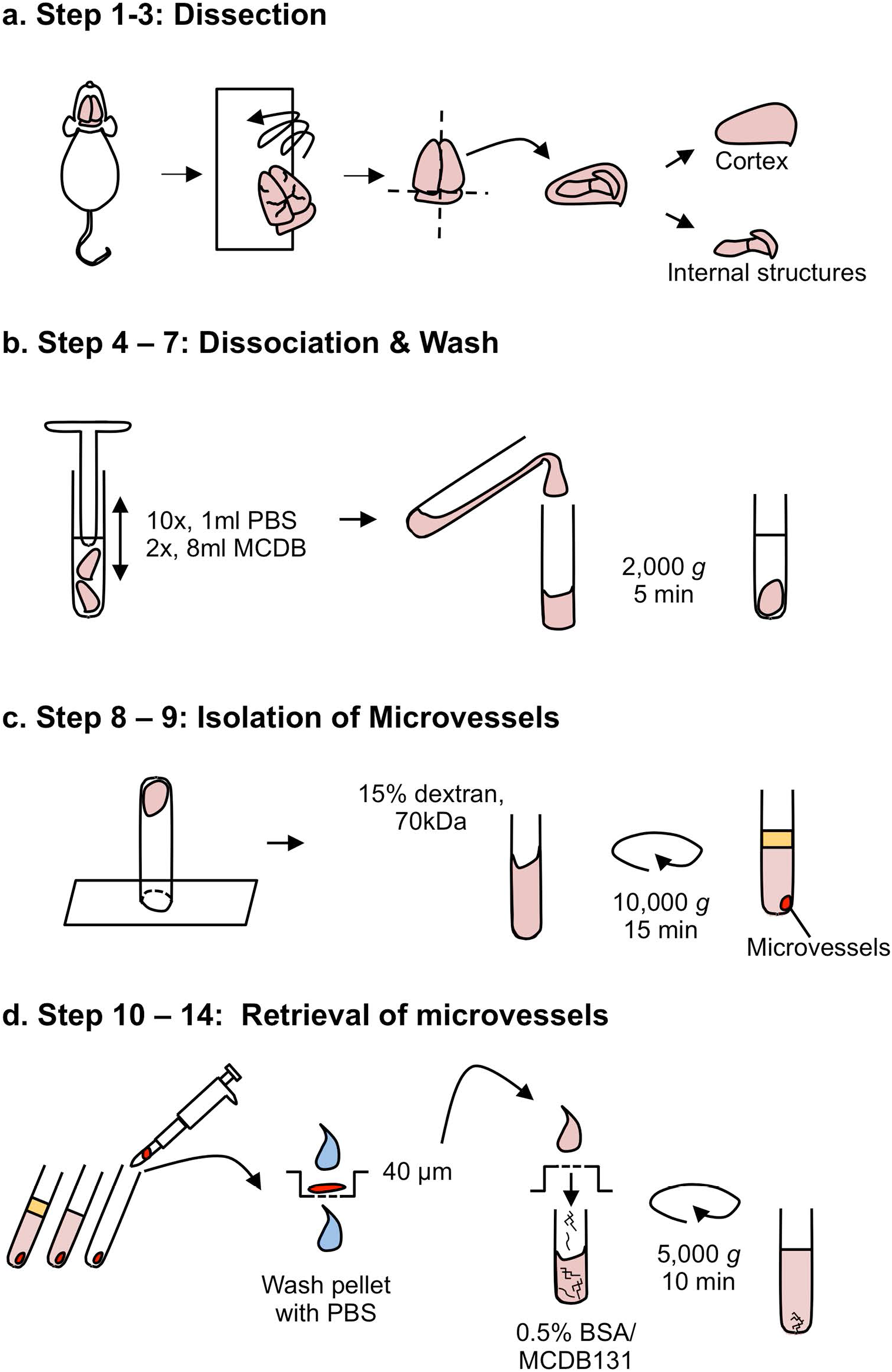
Cortical microvessel isolation protocol overview. **(a)** Dissection. After mouse euthanasia, all steps are conducted in the cold room. Meninges are removed by rolling the brains on blotting paper and cortices are dissected in cold PBS. **(b)** Dissociation of tissue and wash. Cortices are homogenized using a loose fit Dounce grinder and centrifuged (4°C) for 5 min. at 2,000 g. **(c)** Separation of the microvessel fraction by gradient centrifugation. Microvessel pellet is resuspended in 15% 70kDa dextran solution and centrifuged (4°C) for 15 min. at 10,000g. **(d)** Retrieval of microvessels. The top layer containing myelin and brain parenchymal cells is removed. Microvessels are retrieved by pipetting and transferred to a 40μm cell strainer. After being washed with cold PBS, microvessels are collected in a tube by inverting the filter and adding 10 mL of MCDB131 containing 0.5% endotoxin-, fatty acid- and protease free BSA. The suspension is centrifuged (4°C) for 10 min. at 5,000 g to pellet microvessels for further applications.

**Figure 2.**
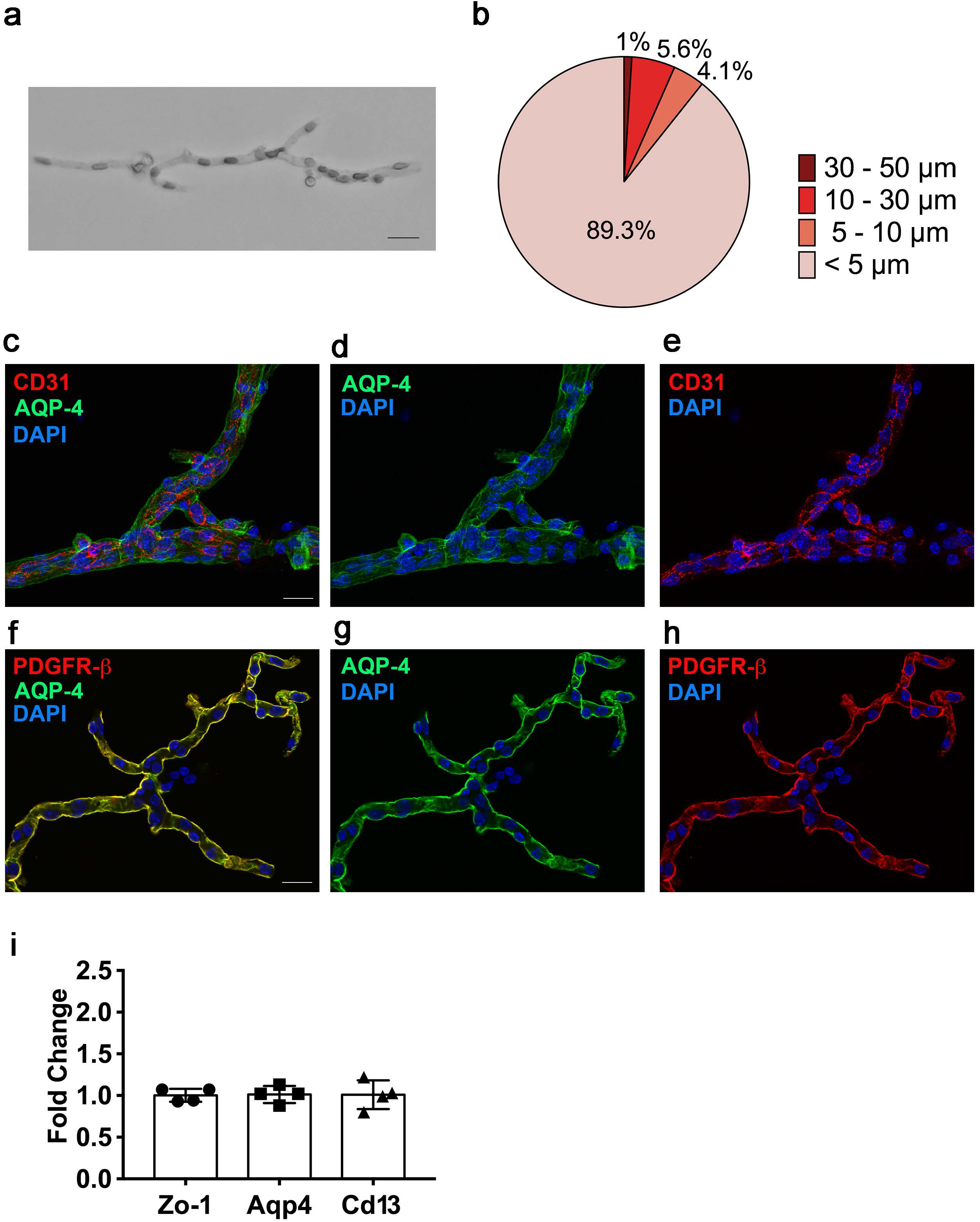
Characterization of microvessel fragments. Isolated microvessels were plated on microscope slides. **(a)** Bright field image of representative microvessels (Scale bar: 20 μm). **(b)** Microvessel diameter was measured under inverted microscope (Olympus CKX41) using the built-in tools of CellSens Entry software (Olympus). 700 microvessels were measured. **(c-h)** Immnuofluorescence analysis of isolated microvessels stained with antibodies for each cell components, **(c, e)** CD31 (endothelial marker, red channel), **(c, d, f, g)** AQP-4 (astrocyte end foot marker, green channel), **(f, h)** PDGFRβ (pericyte marker, red channel), **(c-h)** DAPI (nuclear stain, blue channel). Z-stack images were taken using confocal laser scanning microscope (Olympus, model. FV10i) and processed using Olympus Fluoview ver. 3.0 software. Scale bar indicates 20 μm. (i) The cellular compositions between microvessel preparations were tested. The content of endothelial cells, astrocytes and pericytes in four independent microvessel preparations was assessed by quantification of Zonula occludens-1 (Zo-1, endothelial marker), Aquaporin 4 (Aqp4, astrocyte end foot marker) and Cd13 (pericyte marker) mRNA levels by RT-qPCR. Notice the consistent cell composition between different preparations. RIN ≥9.8. The individual values and the average ±SEM are shown.

**Figure 3.**
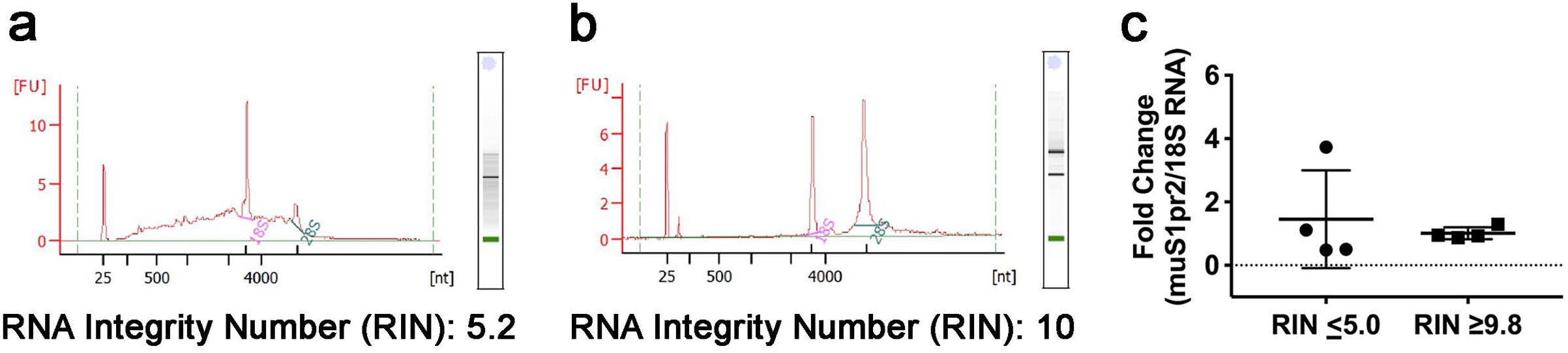
Impact of the microvessel isolation method on the RNA integrity and sample-to-sample variation. Isolation of microvessels was performed in two different environmental conditions. RNA was isolated from microvessels and RNA integrity was determined using the BioAnalyzer 2100 **(a, b)** Bioanalyzer profiles of RNA isolated from microvessels when the preparation was conducted **(a)** at room temperature but all reagents, tools and equipment were kept on ice or cold, **(b)** at 4 °C in a cold room. FU, fluorescence unit, nt, nucleotides. **(c)** Sphingosine-1-phosphate receptor 2 (S1pr2) mRNA levels were quantified by reverse transcription followed by quantitative PCR (RT-qPCR) analysis and normalized by 18S RNA. The individual values and the average ±SEM are shown, n = 4. The coefficient of variation between preparations was 106.2% when RIN ≤ 5 and 18.8% when RIN ≥9.8.

**Figure 4.**
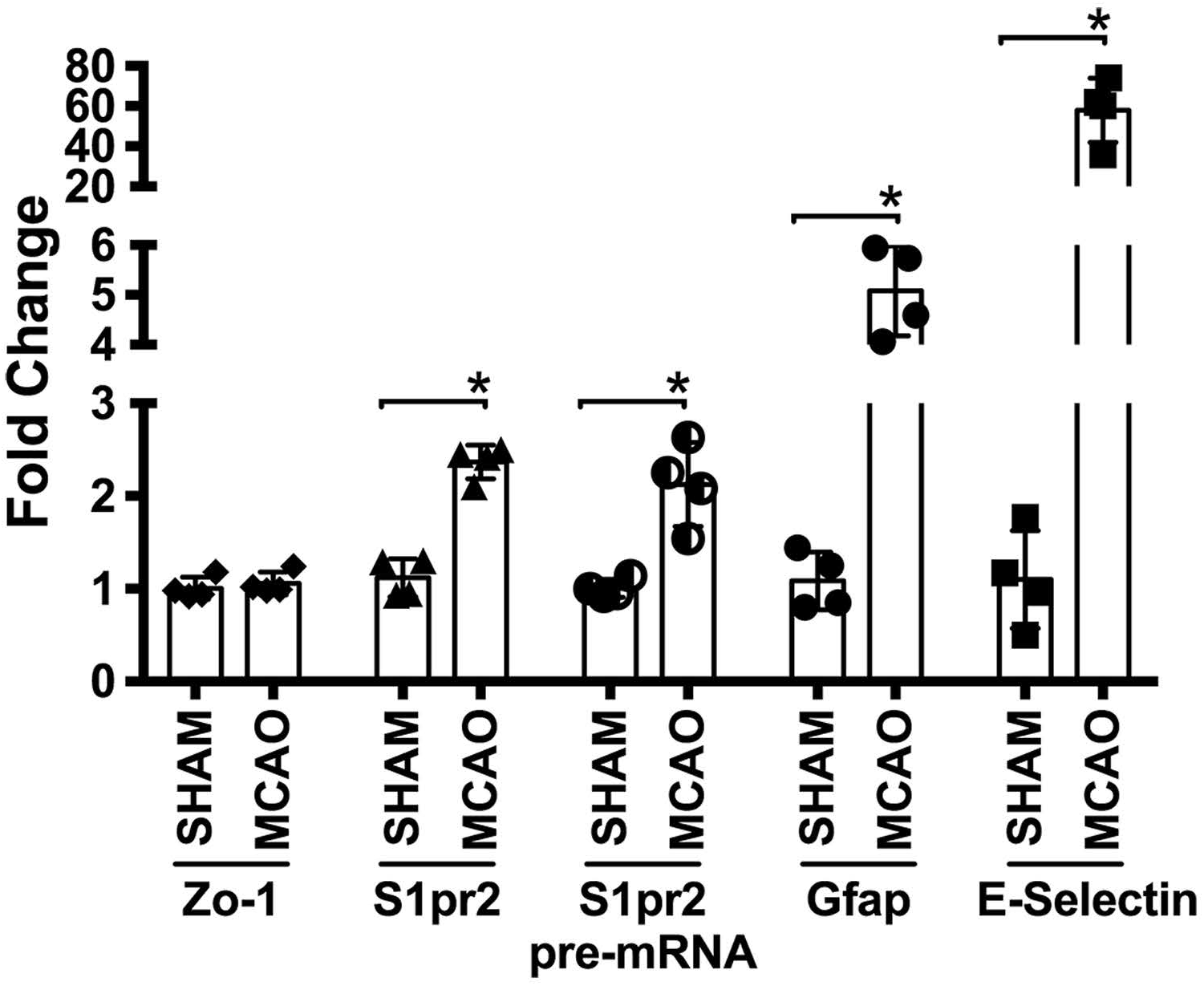
Quantification of changes in gene expression in cerebral microvessels after stroke. Mice were subjected to sham surgery or transient middle cerebral artery occlusion (tMCAO, 45 minutes occlusion followed by reperfusion). 24 hours later, microvessels were isolated from the ischemic (ipsilateral) cortex (MCAO) or sham. Changes in Zo-1, S1pr2, glial fibrillary acidic protein, E-Selectin mRNA (Zo-1, S1pr2, Gfap and E-Selectin, respectively), as well as S1pr2 pre-mRNA levels (S1pr2 pre-mRNA) were quantified by RT-qPCR and normalized by 18S RNA. RIN ≥ 9.8. Fold induction MCAO *versus* sham is shown. The individual values and the average ±SEM are shown. *p <0.05 (t-test). n = 4. The use of laboratory animals was approved by Weill Cornell Medicine Institutional Animal Care and Use Committee.

**Figure 5.**
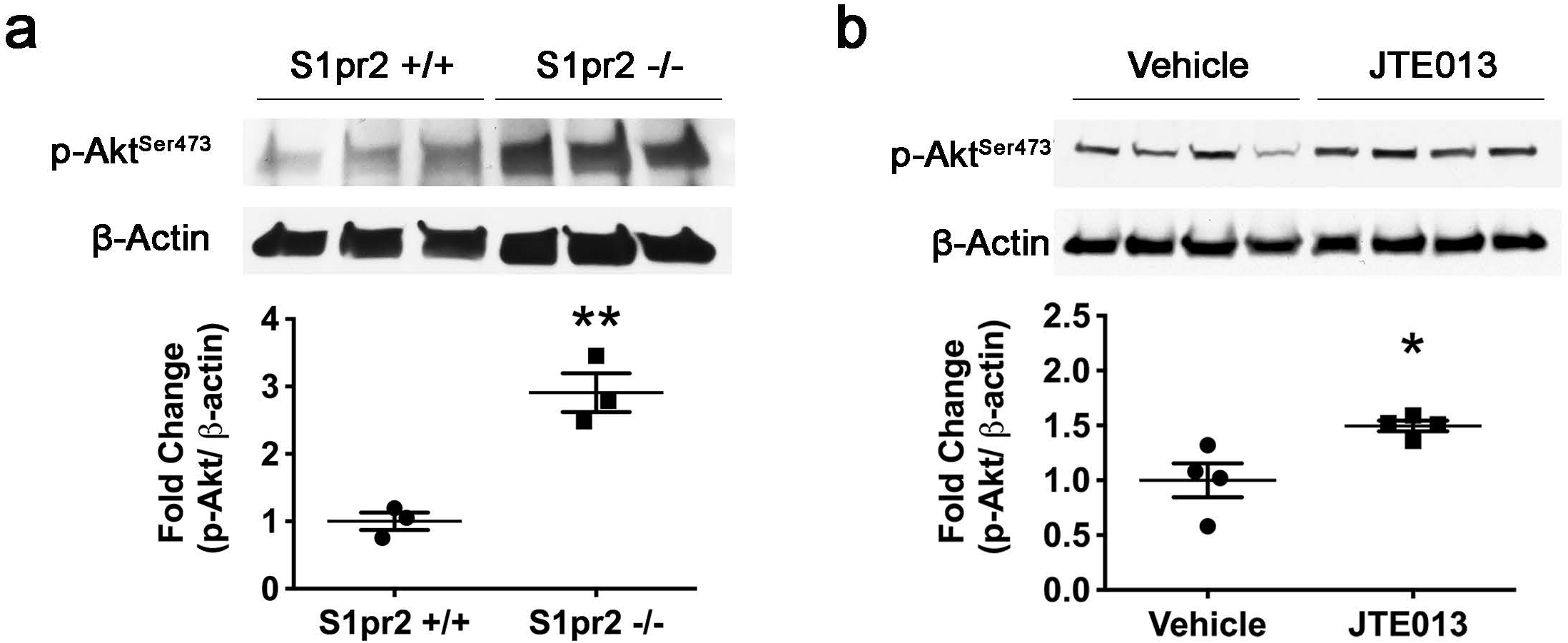
Quantification of phospho-Ser 473 Akt levels in cerebral microvessels in wild type mice after JTE-013 administration. (a) Microvessels were isolated from 8 week-old adult S1pr2 knockout (S1pr2 −/−) mice or wild-type littermates (S1pr2 +/+), and western blot analysis for phosphor-Ser 473 Akt was conducted. n=3. (b) Mice were treated with vehicle or the S1PR2 antagonist, JTE-013 (30 mg kg^−1^) by gavage. 6 hours later, microvessels were isolated and proteins were subjected to Western blot analysis. n=4. Optical density of each lane was measured using the Image J software. Fold induction of p-Akt (normalized by β-actin) in (a) S1pr2−/− *versus* wild type mice or (b) JTE-013-treated *versus* vehicle-treated mice are shown. The individual values and the average ±SEM are shown. *p <0.05 (t-test). The use of laboratory animals was approved by Weill Cornell Medicine Institutional Animal Care and Use Committee.

**Figure 6.**
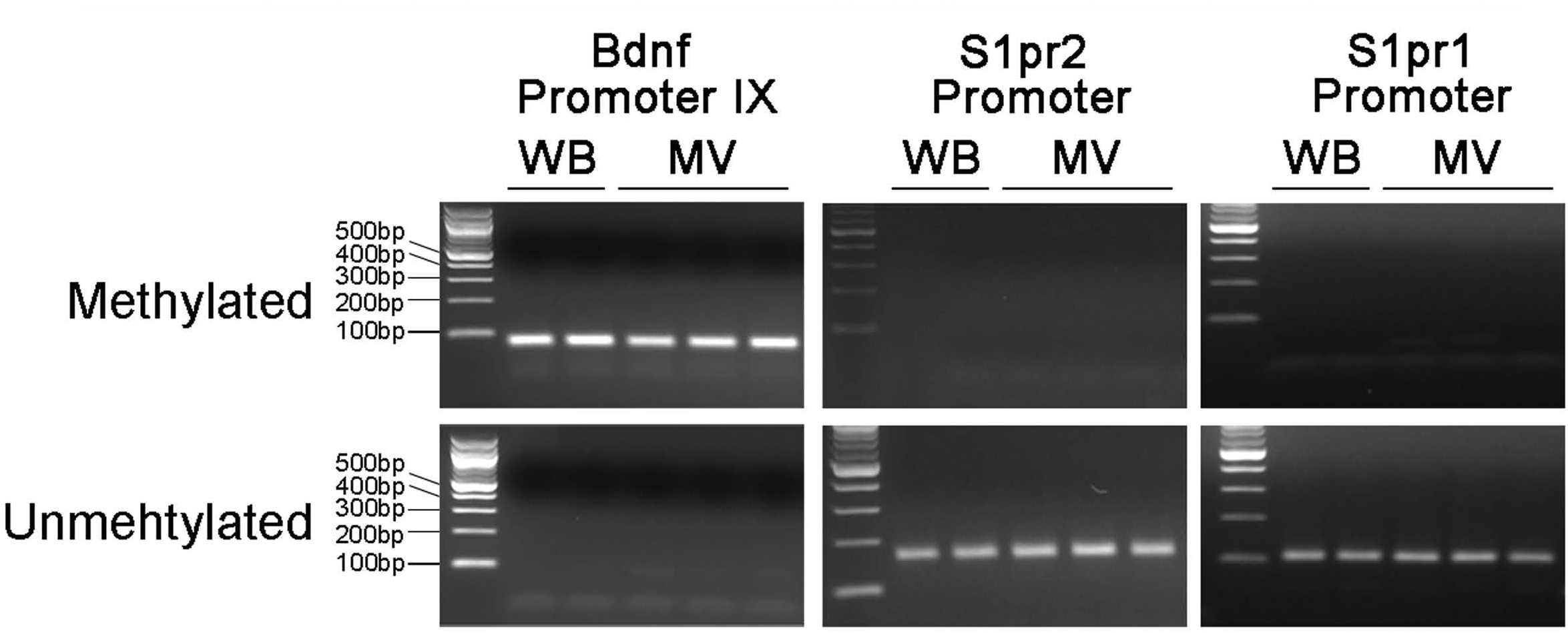
Detection of DNA methylation in Bdnf, S1pr2 and S1pr1 promoter regions in microvessels (MV) and whole brain (WB). Methylation status was examined using methylation specific PCR with specific primers located in Bdnf promoter IX and S1pr1 and S1pr2 core promoter regions. Genomic DNA was isolated from whole brain and microvessel fraction of adult brain (2 month old, C57BL/6J) and subjected to bisulfite treatment. PCR products were analyzed on a 2 % agarose gel. Left lane:100 bp DNA ladder

**Figure 7.**
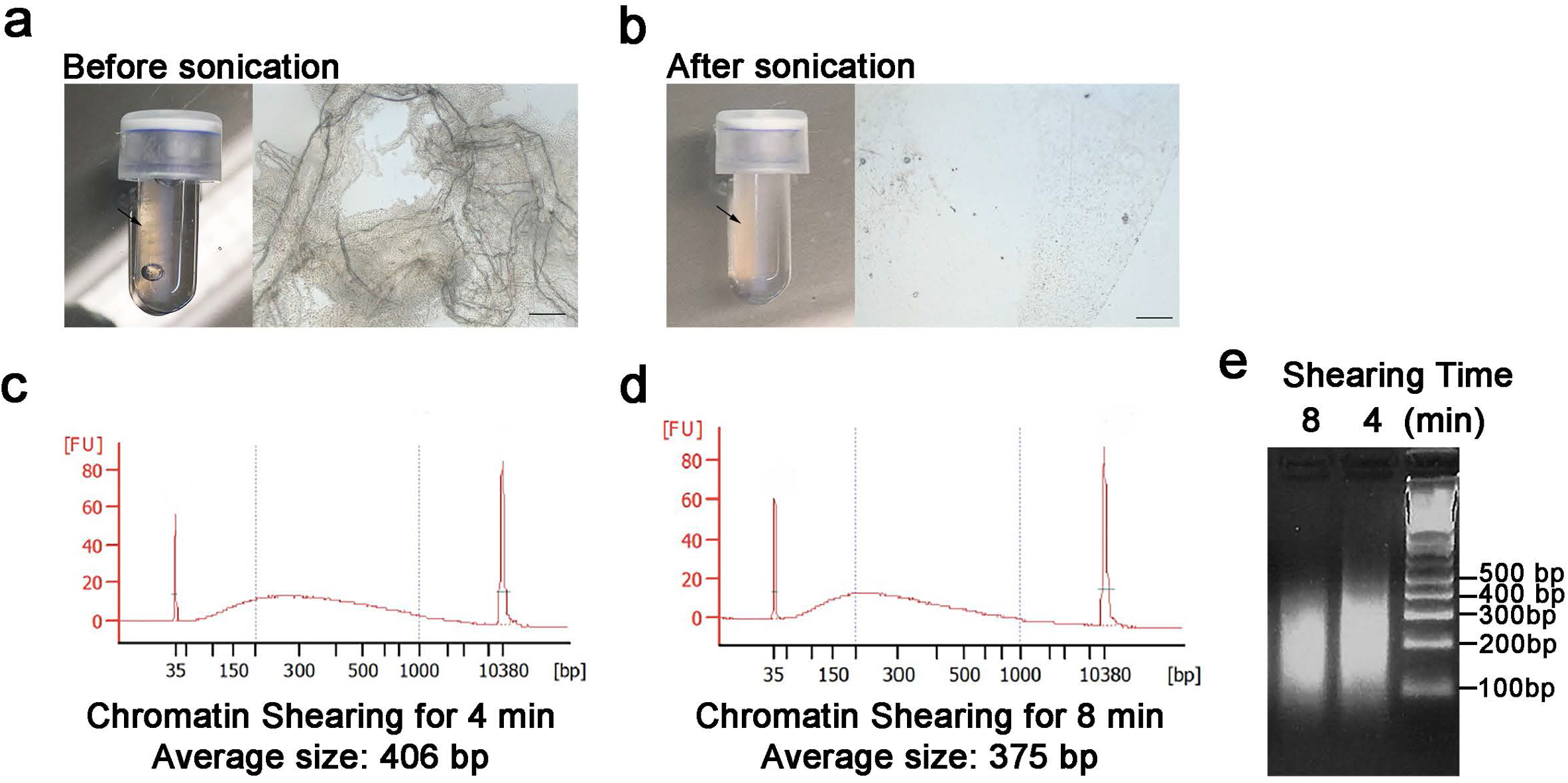
Optimization of chromatin extraction and shearing from cortical microvessels. Isolated microvessels were cross-linked with 1% methanol-free paraformaldehyde and subjected to chromatin extraction protocol. **(a)** Images of lysate after vortexing for 15 minutes in lysis buffer B, as *per* manufacturer’s instructions. Left panel: Arrow indicates visible microvessels in microtube. Right panel: microscopic image of the lysate was taken under inverted microscope (Olympus CKX41, Scale bar: 100 μm) **(b)** Images of lysate after optimization of the conditions to efficiently disrupt microvessels. Left panel: microvessels are no loner visible. Right panel: microscopic image of the lysate. **(c-e)** Impact of the time of sonication on the size of chromatin fragments. After microvessel disruption, extracted chromatin was sonicated for 4 min **(c, e)** or 8 min **(d, e)**. The size of the sheared chromatin was determined by BioAnalyzer **(c,d)** and electrophoresis on a 2% agarose gel **(e)**. Right lane: 100bp DNA ladder.

This protocol will be of great use to researchers intending to understand the molecular mechanisms governing cerebrovascular dysfunction in disease models, or the study of the activation of signaling pathways and intercellular interactions in the cerebral microvasculature *in vivo*. Ultimately, these studies will aid the development of novel therapies targeting the endothelium to mitigate the progression and/or exacerbation of neuronal injury in cerebrovascular and neurodegenerative diseases.

### Development of the protocol and comparison with other methods

To isolate cortical microvessels, we initially used the protocol described by Wu et al., ^14^. While the protocol produced consistent samples in terms of microvessel size and cellular composition, it calls for an enzymatic digestion step. Although samples would potentially be suitable for immunostaining and some biochemical analyses, application of exogenous enzymes and incubation at 37°C cause undesirable molecular and metabolic changes and compromise the integrity of RNA (data not shown). On exclusion of the digestion step, samples contained unwanted debris, as assessed from low-power bright-field images, whereby cellular debris and detached cells could easily be observed. In order to increase the purity of the preparation without the use of enzymatic digestion, we employed an additional wash step (Procedure Step 7) and optimized the dissection and homogenization processes. In order to speed up and optimize the dissection process, we found that gently rolling the brains on blotting paper enabled swift and very efficient removal of the meninges compared to the use of forceps. This technique, which was described in an endothelial cell isolation protocol by Ruck et al. ^11^, considerably shortened the dissection process. In addition, we found that consistent tissue homogenization technique is essential in producing samples of similar composition and containing minimal debris. While existing protocols recommended mincing brain tissue into small pieces and using between three (3) and 20 strokes (1) using Dounce homogenizer ^14,16^, we found that 10 strokes without cutting tissue is enough to homogenize cortices and obtain pure microvessel preparations of consistent cellular composition (Figure 2). This modification shortens the time of preparation and therefore, it is critical to preserve microvessel integrity and prevent RNA degradation.

After optimization of our protocol to obtain pure microvessel preparations of consistent cellular compositions, the next critical factor for the molecular characterization and quantification of changes in gene expression is the quality of the RNA. Therefore, we used RNA quality as the ultimate benchmark in reaching our optimized protocol and next, examined the impact of the microvessel isolation method on RNA integrity. We compared the RNA integrity of microvessel preparations conducted in two different conditions. In one condition, every reagent, tools and equipment were kept on ice but the isolation was performed at room temperature, as previously described ^16^. In another condition, all steps (from the dissection of the brain to the final step) were conducted in the cold room. Interestingly, we found dramatic changes in the RNA integrity of the microvessel preparations obtained by following these two experimental protocols. As shown in Figure 3a, RNA from microvessels isolated at room temperature was partially degraded as assessed by the integrity number (RIN, 5.2). In contrast, the RNA quality was dramatically increased and RNA showed no degradation (RIN, 10) when microvessels were isolated in the cold room (Fig. 3b). We then tested how RNA integrity can affect the reliability of results. We chose the Sphingosine-1-phosphate receptor 2 (*S1pr2*) gene, since we have previously shown that S1PR2 is a low abundance transcript in the endothelium (1-5 mRNA copies/cell) ^18^, and therefore particularly difficult to study in terms of generating consistent results if RNA integrity is compromised. cDNAs were synthesized using RNA isolated from microvessel preparations with RIN ≥9.8 or RIN ≤5.0 and the delta Ct value of each sample to average value of the group was calculated (Fig. 3c). We found that the coefficient of variation between the preparations was 18.8% when RIN ≥9.8, and as high as 106.2% when RIN ≤5.0. Altogether these data indicate that the method of microvessel fraction preparation has a great impact on the integrity of the RNA preparation, which is critical for the accurate quantification of low abundance transcripts.

In summary, we found that compared to previously published protocols, the optimizations of the method described here (i.e. dissection and homogenization techniques, the absence of an enzymatic digestion step and the fact that the entire protocol is conducted at 4°C in the cold room) have a significant impact on the purity and the integrity of the preparation, the consistency of the cellular composition and the quality of the RNA, which significantly affects sample-to-sample variation (Fig. 1) and as a result, the ability to detect changes in gene expression in disease models or upon pharmacological treatments. In addition, we provide a detailed protocol for the successful isolation of RNA, protein, genomic DNA and chromatin from cerebral microvessels for their molecular characterization.

### Applications of the method

Our studies have focused on assessing changes in gene expression or phosphorylated signaling protein levels in disease models or after drug administration. We have shown that this protocol can be used to quantitatively detect changes in gene expression in low abundance and high abundance transcripts in cerebral microvessels in a mouse model of ischemic stroke (Fig. 4). The protocol can also be used to detect changes in protein phosphorylation upon drug administration *in vivo*, which allows the investigation of the activation of signaling pathways in the cerebral microvasculature *in vivo* (Fig. 5). In addition, genomic DNA isolated from microvessels using this protocol can be used to detect the DNA methylation status by methylation-specific PCR after bisulfite treatment (Fig. 6). Bisulfite treated DNA can be used for further methylation analysis such as pyrosequencing or large-scale bisulfite sequencing. Finally, we have shown that chromatin can be extracted from microvessels after crosslinking with 1% methanol free paraformaldehyde and sheared to the appropriate size (Fig. 7), and hence, it could be subjected to chromatin immunoprecipitation (ChIP) assay and further analysis such as ChIP-Seq.

In sum, RNA, protein and DNA extracted from microvessels using the described protocol is suitable for use in many different molecular biology and biochemical assays in which the quality of material used is of vital importance in producing reliable and reproducible data.

### Experimental design

Each microvessel isolation procedure should include control and experimental groups conducted simultaneously to minimize any variation between procedures. The protocol described in this article usually takes about 90 – 120 min. when conducted using 4-5 mice. The overall time of the procedure may differ depending on the level of expertise of the researcher. Therefore, we encourage planning number of samples in accordance with what can feasibly be completed within 2 hours, to preserve microvessel structural integrity, and to prevent degradation of RNA or signaling molecules such as phosphorylated proteins. The expected yields from a microvessel preparation from one mouse are approximately 3 μg of RNA, 3 μg of genomic DNA and 160μg of protein.

In addition, every preparation should be examined for 1) purity and structural features such integrity and size distribution, 2) consistency in the cellular composition of endothelial cells, pericytes and astrocytes, and 3) RNA integrity for gene expression analysis.

The purity and structural features of the microvessel preparations can be examined using a bright field microscope. When the procedure is performed successfully, microvessels should retain their structure as shown in Figure 2a, and brain debris, myelin, white matter and single cells should not be observed in the final preparation. In addition, we encourage evaluating the size distribution of isolated microvessels, when first following the protocol. We found that the majority (∼90%) of microvessels isolated as described in this protocol are < 5 μm in diameter, 4.1% between 5-10 μm, 5.6% 10-30 μm, and 1% 30-50 μm (Fig. 2b).

The cellular composition of microvessels can be analyzed using immunofluorescence staining, quantitative real-time PCR and Western blotting. We have found that microvessels isolated using this protocol retain consistent composition, as assessed by immunofluorescence staining using antibodies for specific blood brain barrier cellular components including: endothelial cells (e.g. CD31), pericytes (platelet-derived growth factor receptor-beta, PDFGRβ), and astrocytic end feet (aquaporin 4, AQP4) (Fig. 2c-h). In addition, it is critical to confirm consistency of the cellular composition between preparations by measuring the expression levels of marker genes for these components such as zonula occludens-1 (Zo-1), CD13, and Aqp 4 (i.e. endothelial cell, pericyte and astrocyte markers, respectively), by quantitative RT-PCR. As shown in Figure 2i, we found that 4 different preparations contained similar level of cell components. When microvessel fragments are isolated for quantification of gene expression or activation of signaling pathway analysis, we strongly recommend to confirm the consistency of cellular composition between preparations by RT-qPCR.

Lastly, when microvessels are to be used to determine changes in RNA expression between experimental groups, every RNA sample isolated should be tested for RNA integrity. We have demonstrated that RNAs can be easily degraded during the procedure (Fig. 3); it is a long process relative to routine whole tissue harvesting and pulverization for RNA extraction, which prolongs the time that microvessels are exposed to endogenous RNases. While we found that performing the entire procedure at 4 °C protects RNA degradation very effectively (Fig. 3), we strongly recommend to examine RNA integrity for every sample before proceeding for further analysis. We used a BioAnalyzer to test RNA integrity, as it gives the quantitative values (RIN 1 to RIN 10), and only requires a small amount of RNA for each assay, but alternative methods such as gel electrophoresis may be used if BioAnalyzer is unavailable.

### Level of expertise needed to implement the protocol

Regarding the level of expertise needed to implement this protocol, it is important to consider that the isolation of microvessels requires dissection of the mouse brain. While a microscope is not necessary for the dissection, competent handling of forceps and other appropriate dissecting tools is required to cleanly and swiftly separate the brain from the skull, and to remove the cortex from the deeper structures of the brain. Further to dissection skills, a person who has both some general rodent handling experience, as well as familiarity with basic biological techniques (such as preparation of solutions) and equipment (such as the centrifuge) should be able to conduct the procedure competently with 2-3 practice experiments. As reference, in order to get intact preparations, the entire protocol needs to be completed in 90-120 min.

### Advantages and limitations of this approach

The key advantages of the proposed method for the isolation of cerebral microvessels are the following:

This method permits the quick isolation of cortical microvessel fragments of consistent cell composition without the need of enzymatic digestion, thereby preserving RNA integrity. Therefore, it enables quantification of changes in gene expression (even low abundance transcripts or pre-mRNA) in disease models, upon *in vivo* pharmacological treatments, or genetic deletion in mice.

Due to the high integrity and cellular consistency of the preparations obtained using this method, additional molecular studies can be conducted to gain mechanistic insights into the molecular mechanisms governing these changes in gene expression. We provide detailed protocols for the determination of methylation status of specific gene promoters and chromatin isolation and shearing.

Since protein post-translational modifications are also retained (e.g. phosphorylation), the method can be used for the quantification of the activation of signaling pathways or changes in the subcellular localization of tight junction proteins upon ligand or drug administration *in vivo*. The data obtained can be correlated with *in vivo* functional assays, such as for instance, determination of blood brain barrier function via injection of specific vascular markers ^19^. Compared to alternative approaches such as brain endothelial cell isolation and *in vitro* culture, the proposed method preserves the physiological blood flow conditions and the cellular interactions with mural cells during the treatments.

When mastered, the technique is easy, fast and it yields intact cerebral microvessel fragments with consistent cell composition.

As any other technique, it has limitations, which are critical to keep in mind during the experimental design and at the time of the interpretation of the results. The main limitations of this technique are the following:

Although the preservation of the cellular interactions between the endothelium and mural cells is a critical factor to maintain the phenotype and the molecular signature of the cerebrovascular endothelium, microvessel fragments consist of, mainly, three cellular components (i.e. endothelial cells, pericytes and end feet of astrocytes). For this reason, acquisition of quantitative data is only possible when cell composition is consistent between experimental groups. Therefore, if the disease model alters the cellular composition, it is recommended to change the experimental design (e.g. choosing an earlier time point) to prevent changes in the cellular components. If this is not feasible, alternative methods, such as single cell isolation and sorting need to be conducted.

Like in other organs, the brain vasculature is heterogeneous and with the proposed method, it is only possible to quantify average changes across the microvessel preparation. This approach can be combined with semi-quantitative techniques such as immunofluorescence analysis or *in situ* hybridization to detect changes in discrete regions or specific cell populations. If information such as cell-specific changes in gene expression or the global distribution of transcript levels across cell populations need to be obtained, other approaches such as single cell RNA sequencing need to be used.

## MATERIALS

## REAGENTS

### Mice and drug administration

- Adult C57BL/6J mice (6-week-old to 4 month old) and S1pr2 null mice ^20^. !CAUTION All animal experiments should be conducted in accordance with the relevant institutional guidelines and regulations. Our protocol for the use of laboratory animals was approved by Weill Cornell Medicine Institutional Animal Care and Use Committee.
- JTE013 (Cayman Chemical. cat. no. 10009458)
- (2-Hydroxypropyl)-β-cyclodextrin (Sigma-Aldrich, cat. no. C0926)

### Microvessels isolation

- 10x PBS (Thermo Fisher, cat.no. 14190250)
- Dextran, molecular weight (MW) ∼70,000 Da (Sigma-Aldrich, cat. no. 31390)
- MCDB 131 Medium, no glutamine (Thermo Fisher, cat. no. 10372019)
- Falcon™ Cell Strainer (Corning Inc. cat. no. 352340)
- Fatty acid free BSA, protease-nuclease free (Millipore Sigma, cat.no.126609)
- LiberaseTM (Roche, cat.no. 5401135001)

### Immunofluorescence staining

- Paraformaldehyde (Sigma Aldrich. cat. no. P6148) !CAUTION Paraformaldehyde is classified as a known cancer hazard. It should be handled only in a working chemical fume hood with personal protective equipment (PPE) such as lab coats, gloves and face shields.
- 1N NaOH (Fisher Scientific. cat. no. SS266-1)
- 1N HCl (Fisher Scientific. cat. no. SA48-1)
- Bovine serum albumin (Fisher Scientific. cat. no. BP-1605)
- Anti-CD31(Dianova, cat.no. DIA-310)
- Anti-Aquaporin 4 (Sigma-Aldrich. cat.no. A5871)
- Anti-PDGFRb (R&D system, cat.no. AF1042-SP)
- DAPI (Sigma-Aldrich, cat.no. D9542)
- VECTASHIELD Antifade Mounting Medium (Vector Labs. cat.no. H1000)

### RNA isolation

- Diethyl pyrocarbonate (Sigma-Aldrich, cat. no. D5758)
- β-mercaptoethanol (Bio-Rad, cat.no. 1610710) !CAUTION β-mercaptoethanol is toxin on inhalation and contact with skin. It should be handled only in a working chemical fume hood with personal protective equipment (PPE) such as lab coats, gloves and face shields.
- Shredder (Qiagen, cat. no. 79654)
- RNase free DNase set (Qiagen, cat.no. 79254)
- RNeasy Mini Kit (Qiagen. cat. no. 74106)

### cDNA synthesis and quantitative PCR

- Verso cDNA (Thermo Fisher, cat.no. AB1453B)
- PerfeCTa SYBR Green Fast Mix, Low ROX (Quanta Bio, cat.no. 95074)
- 1M Tris-HCl, pH 8.0 (Boston BioProduct. cat.no. BM-320)
- 0.5M EDTA, pH 8.0 (Boston BioProduct. cat.no. BM-150)

### Western Blot

- BCA kit (Pierce, cat. no. 23227)
- Protease Inhibitor Cocktail Set I (CalBiochem, cat. no. 539131)
- Spectra^™^ Multicolor Broad Range Protein Ladder (Thermo Fisher cat. no. 26634)
- 1M HEPES Buffer, pH 7.3 (Fisher Scientific. cat. no. BP299100)
- Triton X-100 Detergent (Bio-Rad. cat. no. 1610407)
- Sodium deoxycholate (Sigma-Aldrich. cat. no. D6750)
- Sodium dodecyl sulfate (Fisher Scientific. cat. no. BP166-100)
- 5M Sodium chloride (Promega Corporation. cat. no. V4221)
- Magnesium chloride solution, 1M (Sigma-Aldrich. cat. no. M1028)
- β-glycerophosphate disodium salt hydrate (Sigma-Aldrich. cat. no. G6251)
- Sodium orthovanadate (Sigma-Aldrich. cat. no. S6508)
- Sodium fluoride (Sigma-Aldrich. cat. no. 215309)
- 4-15% Precast Protein Gels (Bio-Rad. cat. no. 4561084)
- 10x Tris/Glycine Buffer for Western Blots (Bio-Rad. cat.no.1610771)
- 10x Tris/Glycine/SDS (Bio-Rad. cat.no.1610772)
- TBS buffer (Bio-Rad. cat.no.170-6435)
- Tween20 (Fisher Scientific. cat. no. BP337-100)
- Immobilon-P Membrane, PVDF, 0.45 μm (Millipore. cat. no. IPVH00010)
- ECL Western Blotting Substrate (Pierce. cat. no. 32106)
- Blue Autography film (Crystalgen. cat. no. CGFB-507)
- Monoclonal Anti-β-Actin antibody (Sigma-Aldrich. cat. no. A5316)
- Phospho-Akt (Ser473) (193H12) Rabbit mAb (Cell Signaling Technology. cat. no. 4058)
- Restore^™^ Western Blot Stripping Buffer (Pierce. cat. no. 21059) Genomic DNA isolation
- DNA Blood & Tissue Kit (Qiagen. cat. no. 69504) Methylation Specific PCR
- EpiTeck Bisulfite kit (Qiagen, cat.no. 59104)
- EpiTeck MSP kit (Qiagen, cat.no. 59305)
- Agarose (Denville Scientific Inc. cat. no. GR140)
- Ethidium Bromide Solution (Bio-Rad, cat. no. 1610433)
- 100bp DNA ladder (New England BioLabs Inc, cat. no. N3231)

### Chromatin Shearing

- RNase A (Sigma-Aldrich, cat.no. R4642)
- Proteinase K (Roche, cat. no. 50-100-3393)
- truChIP Chromatin Shearing Kit with Formaldehyde (Covaris. cat. no. 520155)
- PCR purification kit (Qiagen, cat. no. 28104)

### EQUIPMENT

- Dounce tissue grinder set (Sigma-Aldrich. cat. no. D9063)
- 14 mL round bottom tubes (Nunc^™^. cat. no. 150268)
- Ultracentrifuge (SORVALL, model RC-5C Plus, Rotor: 05-SS-34)
- Microcentrifuge tube
- Surgical fine forceps (Fine Science Tools. cat. no. 11271-30)
- Petridish
- Inverted light microscope (Olympus. model CKX41)
- Freezer
- Shaker
- Rotator
- Fume hood
- Superfrost Microscope slides (Fisher Scientific. cat. no. 12-550-123)
- Cover glasses (Fisher Scientific. cat. no. 12-544-12)
- Confocal laser scanning microscope (Olympus, model. FV10i)
- Real-time PCR system (Applied Biosystems. model. ABI 7500 Fast)
- Pulsing Vortex Mixer (Fisher Scientific. model. 02-215-375)
- BioAnalyzer 2100 (Agilent)
- Sonicator (Fisherbrand^™^ Q55 Sonicator. model FB50)
- Gel documentation system (Bio-Rad)
- Covaris M220
- microTUBE Sap-Cap AFA Fiber (Covaris)

### REAGENT SETUP

**S1pr2 null mice** Mice carrying targeted description of the S1pr2 gene are maintained on a mixed mixed C57BL/6;129Sv genetic background (five times backcrossed to C57BL/6) ^20^. Wild-type littermates should be used as controls for experiment on S1pr2 knockout (KO) mice.

**JTE013 stock solution** Dissolve JTE013 in DMSO at a final concentration of 100 mM. Make sure that it completely goes into solution. Aliquots can be stored up to 6 months at −20 °C.

**JTE013 gavage administration** Prepare a solution of 2% (2-Hydroxypropyl)-β-cyclodextrin in PBS/saline/H_2_O. Filter to sterilize and aliquot to keep it sterile. Just before gavage, prepare a suspension of JTE013 at 3 mg/ml in 2 % (2-Hydroxypropyl)-β-cyclodextrin solution. Vortex right before gavage to homogenize the suspension.

**15 % Dextran/PBS (wt/vol)** Weigh1.5 g of dextran in a 50 ml conical tube and add 8 ml of PBS. Rotate the tube until it completely dissolves at 4°C. It takes about 30 mins. Adjust final volume to 10 ml. Solution can be stored for 1 week at 4 °C.

**0.5 % BSA/MCDB13 (wt/vol)** Dissolve 0.5 g of fatty acid-protease-endotoxin free BSA in 10 ml of 1x PBS using a 15 ml conical tube to make 5 % BSA/PBS solution. Once solution looks clear, add 1 ml of 5 % BSA/PBS to 9 ml of MCDB13 medium. The 5 % BSA/PBS solution can be stored for up to 6 months at −20 °C.

**4 % PFA/PBS (wt/vol)** Weigh 20 g of paraformaldehyde and add to pre-heated Milli-Q water at 60 °C in a glass beaker. Cover and maintain at 60 °C. Add 2 N NaOH until solution gets clear, remove the beaker from the heat and add 50 ml of 10x PBS. Adjust pH to 7.2 using HCl and adjust final volume to 500 ml. Sterile filter with 0.2 μm bottle top filter and aliquot to small volume and store up to 3 months at −20 °C !CAUTION Paraformaldehyde is classified as a known cancer hazard. It should be handled only in a working chemical fume hood with personal protective equipment (PPE) such as lab coats, gloves and face shields.

**0.1 % NP/40 PBS (vol/vol)** Add 1 ml of NP-40 to 999 ml of 1x PBS in a plastic beaker and mix well using a magnetic stirrer. Sterile filter using 0.2 μm bottle top filter and store at room temperature protected from light for up to 6 month.

**Blocking Solution for immunofluorescence staining (wt/vol)** Dissolve 5 g of BSA in 100 ml of 1x PBS and mix well. Once solution looks clear, sterile filter and make 10 ml aliquots. This solution can be stored for up to 6 months at −20 °C.

**DEPC water** Add 1 ml of DEPC to 1L of Milli-Q water in a clean glass bottle and mix well and incubate for 3 hr at R.T with occasional swirling. Autoclave for 15 mins and store at room temperature for up to 12 months.

**1X TE buffer** Add 5 ml of 1M Tris-HCl, pH 8.0 and 1ml of 0.5 M EDTA, pH 8.0 to 500 ml of Milli Q water and mix well. Autoclave for 15 mins and store at room temperature for up to 12 months.

**1M Na_3_VO_4_** Weigh 1.839 g and dissolve in 10 ml of Milli-Q water. Aliquots in 500 μl and store up to 12 months at −20 °C. Avoid free and thaw.

**1M NaF** Weigh 0.42 g and dissolve in 10 ml of Milli-Q water. Aliquots in 500 μl and store up to 12 months at −20 °C. Avoid free and thaw.

### EQUIPMENT SETUP

**Ultracentrifuge** Pre-chill it to 4 °C before use.

**Covaris M220** Fill water to the chamber according to manufacturer’s instruction and prechill before use it.

### PROCEDURE

#### Microvessels Isolation • Timing 1-2 h

**♦ CRITICAL STEP** The entire procedure should be carried out in a cold room.

1. Sacrifice mice in CO**2** chamber for 2-3 mins, perform cervical dislocation and bring the mice immediately to cold room. Use at least one control and one treated mouse (or equivalent) per procedure.
2. Carefully remove brains and transfer immediately to a petri dish containing MCDB 131 medium and rinse in the medium. Ensure entire brain remains submerged in the medium. **♦ CRITICAL STEP** RNAs get degraded as soon as mouse is euthanized. Perform step 1 and 2 on one mouse at one time and keep the brain in the cold MCDB 131 medium on ice until entire group is processed. We encourage having 4-5 mice per isolation because increased number of mice can slow down entire process.
3. Dissect the brain:

a. Using closed forceps, gently and evenly roll the brain on blotting paper to remove meninges and meningeal vessels. Place brain in PBS (which provides greater visibility than MCDB 131 medium for optimal dissection).
b. Using a razor blade, slice brain sagittally and remove cerebellum.
c. Isolate cortices by removing all deep brain structures including the hippocampus, and residual white matter, being careful to avoid damaging the cortices. While as much of the above should be removed as possible, a certain amount of white matter is likely to remain attached. Since this will be removed in future steps, do not attempt to remove all white matter tissue at the cost of damaging the cortices. ? TROUBLESHOOTING
d. Place isolated cortices to MCDB 131 medium or proceed to step 4.
4. Homogenize cortical tissue using a loose fit, 7 ml Dounce tissue grinder. Perform an initial 10 strokes in 1 ml MCDB 131 medium at a steady pace, without twisting the pestle.
5. Add 7 ml of MCDB131 medium to homogenate from step 4 and perform additional 2 strokes in a total of 8 ml.
6. Pour homogenized tissue into a 14 ml round-bottom centrifuge tube. Take care to alter the volumes using additional MCDB 131 medium, so that each sample is evenly balanced.
7. Centrifuge at 2,000 *g* for 5 mins, 4 °C. Pour out supernatant and rest the inverted tube on a paper towel, to absorb excess medium/debris clinging to the sides. **♦ CRITICAL** STEP This is a wash step, and helps with the removal of small pieces of white matter debris that have remained attached to the sample.
8. The pellet must be resuspended in 15 % dextran/PBS.

I. In order to dissociate the pellet evenly, add an initial 1 ml dextran solution to the pellet, and use it to sluice the pellet gently from the side of the tube.
II. Resuspend the floating pellet with 5-6 steady uses of the pipet, taking up the entire pellet with each use.
III. Add a further 7 ml dextran solution and balance with additional dextran solution.
9. Centrifuge at 10,000 *g* for 15 mins, 4°C. The red, microvessel-containing pellet will appear. **♦ CRITICAL STEP** If pellet does not appear to be in clean red color, it indicates contamination with white matter, myelin or debris. ? TROUBLESHOOTING
10. The supernatant should be removed carefully to avoid contaminating the pellet with debris. Begin by pipetting off the top, brain tissue-containing layer completely, then aspirating the underlying solution, which will contain a small amount of debris. ? TROUBLESHOOTING
11. Remove the pellet using 1 ml PBS. Sluice the pellet from the side of the tube, and take up into the pipet tip with the absolute minimum of PBS that has passed into the tube. **♦ CRITICAL STEP** Using more than 1 ml of PBS is discouraged. You will see some of debris on the wall of tube that can be easily taken by pipetting. Using small volume is beneficial to minimize the contamination.
12. Transfer the pellet onto a 40 μm cell strainer, and wash through with <10 ml PBS. This step is intended to remove debris that has remained slightly adhered to the microvessels as well as loose debris. ? TROUBLESHOOTING
13. Reverse the filter and retrieve the microvessels using 10 – 20 ml of MCDB 131 medium containing 0.5% fatty-acid, protease-, and endotoxin free BSA. ***Bright field and phase contrast Image***

> To quickly evaluate purity of microvessels by microscopic method, take 50 ul of microvessels suspension from step 13, and plate onto a 35 mm culture dish for 5 mins at room temperature. Once microvessels attaché to the surface of the dish, observe under inverted microscope.
14. Centrifuge at 5,000 *g* for 10 mins, 4°C. This will yield the final microvessel pellet. ? TROUBLESHOOTING ? TROUBLESHOOTING Troubleshooting advice can be found in Table 2.

**• Timing**

#### Microvessels isolation (90 – 120 min)

Steps 1-3, euthanasia and dissection of the brain: 5-7 min per mouse
Steps 4-7, homogenization cortices and centrifugation: 20 min
Step 8, removal of supernatant and re-suspending pellet in dextran: 10-15 min
Step 9-13, centrifugation and retrieval of microvessels pellet: 30-40 min
Step 14, final centrifugation and resuspend in proper lysis buffer: 15-20 min

#### Analysis of microvessels

##### A. Immunofluorescence characterization • Timing 24 h

i. Add 8 ml of 4% PFA/PBS into a 35 mm petridish.
ii. Submerge the cell strainer containing microvessels from step 12 into 4 % PFA/PBS and fix microvessels for 15 mins with gentle shaking.
iii. Place the cell strainer on a 50 ml tube and wash through with 10 ml of PBS.
iv. Reverse filter on another 50 ml tube, and collect microvessels by thoroughly applying 15 −20 ml of 1% BSA/PBS onto the surface of the filter.
v. Centrifuge the suspension at 2,000 *g* for 10 mins, 4 °C.
vi. Aspirate and discard the 1 % BSA/PBS and resuspend pellet in 400 μl of PBS.
vii. Drop 50 μl of suspension onto a Superfrost microscope slides and air dry slides. It takes about 30 mins. **￭ Pause Point** Slides can be stored at −80 °C for up to 12 months before processing.
viii. Wet the microvessels with 1x PBS for 10 mins.
ix. Permeablize the microvessels with 0.1 % NP40/PBS for 15 mins.
x. Block the microvessels with 5% BSA/PBS for 1 hr.
xi. Incubate the microvessels overnight at 4°C with the primary antibodies CD31 (1:200), PGDFRβ (1:200), AQP4 (1:500) diluted in 5 % BSA/PBS.
xii. After three washes with 0.1 % NP40/PBS, incubate the microvessels for 1 hr at room temperature in the dark with appropriate secondary antibodies conjugated with Alex488 or Alex568.
xiii. Wash three times with 0.1 % NP40/PBS.
xiv. Counter stain the microvessels with DAPI for 1 mins.
xv. Wash once with 0.1 % NP40/PBS.
xvi. Air dry the slide and mount with fluorescence mounting medium. Protect slides from the light and store at 4 °C.
xvii. Observe staining under confocal microscope.

##### B. RNA isolation • Timing 2-3 h

i. After centrifugation from step 14, discard supernatant and quickly remove residual medium by inverting the tube on a paper towel.
ii. Add 700 μl of RLT buffer provided from RNeasy Kit (Qiagen).
iii. Resuspend microvessels pellet in RLT buffer with 20 – 25 times of pipetting and vigorous vortexing for approximately 30 sec to completely disrupt microvessels.
iv. Snap freeze in liquid nitrogen and store at −80 °C up to 6 months. **￭ Pause Point** Lysate can be stored at −80 °C before processing
v. Isolate RNA according to instruction from Qiagen and include DNaseI treatment step as described in manufacturer’s instruction.
vi. Measure RNA integrity by BioAnalyzer. **♦ CRITICAL STEP** RNA integrity should be tested prior to downstream analysis of mRNA.

##### C. Protein extraction • Timing 2 h

i. After centrifuge from step14 discard supernatant by converting the tube containing pelleted microvessels and stand on paper towel for a few second.
ii. Add 80 μl of ice-cold protein lysis buffer (50 mM HEPES pH 7.5; 1% Triton; 0.5% sodium deoxycholate; 0.1 % SDS; 500 mM NaCl; 10 mM MgCl_2_; 50 mM β-glycerophosphate; 1x Protease inhibitor cocktail; 1 mM Na_3_VO_4_; 1 mM NaF) **♦ CRITICAL STEP** Add protease inhibitor cocktails and phosphatase inhibitors (Na_3_VO4 and NaF) to protein lysis buffer fresh before use.
iii. Sonicate pelleted microvessels for 3 × 5 seconds at 50 amplitude using a probe type sonicator on ice.
iv. Vortex lysate under maximum speed for 15 mins, 4 °C. This step will help lyse microvessels adequately. **♦ CRITICAL STEP** Keep the tip of probe to avoid foaming during sonication.
v. Centrifuge lysate at 10,000 *g* for 10 mins, 4 °C and transfer supernatant into a new tube. **￭ Pause Point** Protein can be stored at −80 °C up to 6 months before processing.
vi. Measure protein concentration using bicinchoninic acid (BCA) assay according to manufacturer’s instruction. We normally obtain 150-160 μg of protein from one brain.

##### D. Crosslinking and chromatin shearing • Timing 24 h

Note: We used a commercially available kit, truChIP^™^ Chromatin Shearing Reagent Kit from Covaris. Here we describe only steps optimized from the manufacturer’s instruction for usage of microvessels.

i. Add 8 ml of 1 % methanol free PFA solution provided from the kit into a 35mm petridish. **!CAUTION** Paraformaldehyde is classified as a known cancer hazard. It should be handled only in a working chemical fume hood with personal protective equipment (PPE) such as lab coats, gloves and face shields.
ii. Transfer the cell strainer containing washed microvessels from step 12 to petridish and incubate for 5 mins at room temperature with gentle shaking. **♦ CRITICAL STEP** Make sure the strainer submerged completely into 1% methanol free PFA solution during the crosslinking.
iii. Add quenching buffer provided from the kit as described in manufacturer’s instruction to stop crosslinking.
iv. After a brief wash with 2ml of PBS, retrieve fixed microvessels from the strainer with 10 ml of 1% BSA/PBS, and centrifuge at 2,000 *g* for 10 mins.
v. Discard supernatant and resuspend the pellet in the lysis buffer B provided from the kit. We combine microvessels from two brains in 300 μl of lysis buffer B. **￭ Pause Point** Microvessels pellet can be stored at −80 °C before processing up to 6 month.
vi. Incubate lysate for 15 mins, 4 °C on a rotator and transfer lysate to the microTUBEs.
vii. Set the CovarisM220 at 5 % duty factor, 50 P.I.P, 200 cpb, and sonicate lysates for 3 mins. **♦ CRITICAL STEP** We incorporated a mild sonication steps due to difficulties of pulverizing microvessels. To maximize yield of chromatin extraction, it is critical to disrupt and lyse microvessels properly and completely. After step vii, the structure of microvessels should not be visible.
viii. Centrifuge lysate from step vii at 1,700 *g* for 5 mins, 4 °C.
ix. Add 600 μl of wash buffer provided from the kit and incubate for 5 mins on a rotator, 4 °C, followed by centrifugation at 1,700 *g* for 5 mins to collect nuclear extract.
x. Resuspend pellet in 550 μl of shearing buffer that provided from the kit, and vortex until it is mixed.
xi. Transfer chromatin to the microTUBE and sonicate with the default setting of Low Cell Chromatin Shearing Protocol for M-Series as described in manufacturer’s instruction for 4 or 8 mins.
xii. Process sheared chromatin according to manufacturer’s instruction to check the size of sheared chromatin by electrophoresis using 2 % agarose gel or by BioAnalyzer.

#### ANTICIPATED RESULTS

##### A. Quantification of changes in gene expression

First, we aimed to determine whether the isolated microvessels retained the expected molecular changes in the brain vasculature in a disease model, namely, the transient middle cerebral artery occlusion (tMCAO) mouse model of ischemic stroke. 2-month-old C57/Bl6 mice were subjected to sham surgery or transient focal ischemia (tMCAO) for 45 mins as we previously described ^18^. 24 hours after reperfusion, cortical microvessels were isolated from the ischemic hemisphere (ipsilateral). Ipsilateral cortices from two mice were pooled.

RNA was isolated from microvessels of each group, and levels of gene expression were quantified by reverse transcription and quantitative real time PCR. The primer sequences used for analysis are indicated in Table 1. As shown in Figure 4, consistent with previous reports, the endothelial pro-inflammatory gene, E-Selectin and the astrocyte activation marker, Gfap, were significantly upregulated in cerebral microvessels in the ipsilateral cortex compared to the contralateral (57.9±14.8-fold and 5.1±0.8-fold induction, respectively). We also aimed to investigate if changes in S1pr2 mRNA, one of the receptors for the bioactive lipid sphingosine-1-phosphate (S1P) ^21^, could be detected in cerebral microvessels *in vivo*. We recently showed that S1pr2 expression increases upon ischemia-reperfusion (I/R) injury in mouse brain and *in vitro* in endothelial cells ^18,22^, Consistent with these studies, S1pr2 mRNA expression was increased 2.4±0.2-fold in cortical microvessels upon I/R injury. In addition, we tested the levels of S1PR2 pre-mRNA levels, which is normally less abundant than mature mRNA. We found that S1PR2 pre-mRNA levels were also increased 2.1±0.4-fold in microvessels in response to I/R injury, suggesting that S1PR2 is regulated at the transcriptional level upon I/R injury. The levels of the endothelial tight junction protein zonula occludens 1 (Zo-1), were not changed (Fig. 4), as well as the astrocyte and pericyte markers (aquaporin 4 and Cd13, respectively, not shown), indicating that the cellular composition was similar between preparations.

**Table 1.** Troubleshooting table.

| Step | Problem | Possible reason | Solution |
| --- | --- | --- | --- |
| 3 | Delayed or improper dissection of cortices | Usage of inappropriate tools can cause difficulties | Use curved end shape forceps |
| 9 | Pellet does not seem red or covered with white layer | Pellet was not properly resuspended | Mix pellet from step 7 completely with 15% dextran/PBS by pipetting |
|  | Pellet does not appear at the bottom of tube | Concentration of dextran solution was incorrect | Confirm that dextran completely dissolves. Do not exceed the correct final volume |
| 10 | Debris is too close to the pellet. It is difficult to get clean pellet | Dextran was not mixed thoroughly with brain lysate or used small volume of dextran solution | Increase volume of dextran and mix the pellet thoroughly by pipetting. If necessary, wipe out debris from the wall of tube using Kimwipe |
| 12 | Pellet gets trapped inside the pipet tip | Microvessels can easily adhere to surface of plastic | Pre-wet pipet tip by pipetting in and out repeatedly 1x PBS |
| 14 | Pellet is very small or invisible | Microvessels remain in the cell strainer or BSA is not added to medium | Use enough volume to retrieve microvessels from the strainer. Add BSA (0.5%) to pellet microvessels |

**Table 2.**
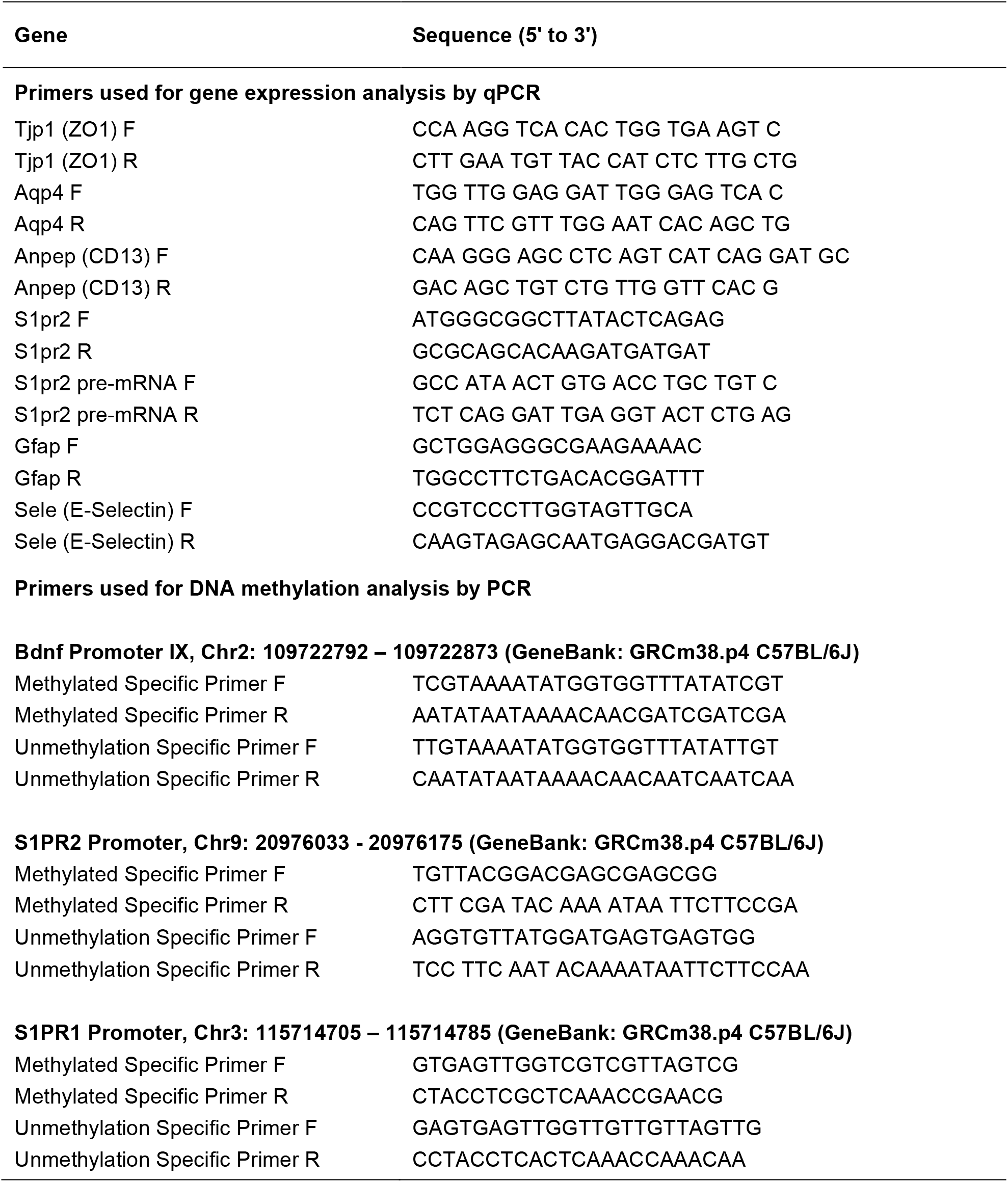
Primer sequences.

##### B. Phospho-protein changes

Given the relative difficulties in measuring changes in protein phosphorylation *in vivo*, we tested if our microvessel isolation protocol was suitable to capture these changes. We and others have previously shown that S1P, which is very abundant in plasma and it is produced and released mainly by endothelial cells, erythrocytes and platelets ^23^, activates S1PR2 in the endothelium giving rise to inhibition of Akt phosphorylation (Ser473) via the Rho-Rho dependent kinase (ROCK)-Phosphatase and tensin homolog (PTEN) pathway ^24-26^. Therefore, we aimed to compare phospho-Ser 473-Akt levels in cortical microvessels of mice bearing genetic deletion of the S1pr2 gene or upon acute inhibition of S1PR2 signaling by administration of the S1PR2 antagonist JTE013. We first compared Akt phosphorylation levels in microvessels isolated from S1pr2 −/− brain cortex with wild type. As shown Figure 5A, we found that phospho-Akt levels were significantly higher in S1pr2 null mice compared to their wild type littermates (2.9±0.29 fold). In addition, phospho-Akt levels were also significantly increased (1.5±0.05 fold) in cerebral microvessels 6 h after administration of JTE013 (30 mg/kg) by gavage. These data are consistent with our previous *in vitro* signaling studies in endothelial cells ^24,27^ and indicate that the method described allows the quantification of changes in protein phosphorylation, and thus, it can be used to conduct signaling studies in the brain microvasculature *in vivo*.

##### C. Assessment of DNA methylation

We also tested if genomic DNA isolated from microvessels can be used to detect epigenetic modifications such as methylation of gene promoters. Primers to detect specific methylated or un-methylated DNA were designed using MethPrimer (http://www.urogene.org/methprimer/) and are shown in Table 1. Genomic DNA (gDNA) isolation, bisulfite treatment, and methylation specific PCR were performed according to manufacturer’s instruction. We first tested promoter IX of brain derived neurotrophic factor (Bdnf) gene since it has been shown to be methylated in adult mouse brain ^28^. We isolated gDNA from microvessels as well as whole brain of adult mice and treated with bisulfite. Then, PCR was conducted with primers to detect methylated or un-methylated DNA (Table 1). As shown in Figure 6, we were able to detect methylation of Bdnf promoter IX while no bands were amplified with the primers to detect unmethylated DNA. These data indicate that Bdnf promoter IX is methylated in both in brain as well as in cerebral microvessels, consistent with previous reports ^28^. We then tested the methylation status of S1pr1 and S1pr2 CpG islands in their core promoters. The information of genomic features including core promoter regions and CpG island was obtained from Ensemble (http://www.ensembl.org) and Genome Browser (https://genome.ucsc.edu). Interestingly, we found that both S1pr1 and S1pr2 CpG islands were unmethylated, which is consistent with the expression of S1PR1 and S1PR2 in the cerebrovascular endothelium. These data indicate that microvessels can serve as a useful material to detect genomic DNA methylation. As described earlier, the bisulfite treated gDNA can be subjected to pyrosequencing to quantify the degree of methylation in CpG islands of the specific gene promoters or to bisulfite whole genome sequencing to discover a broad range of methylation status which should generate valuable information to understand mechanism of regulation in gene expression^29^.

##### D. Optimization of chromatin extraction and shearing from cortical microvessels

ChIP assay is a very valuable technique to determine histone modifications as well as binding of proteins (e.g. RNA polymerases, transcription factors) to specific regions of the chromatin both *in vitro* and *in vivo*. However, when conducting ChIP assays in tissues, chromatin extraction and shearing to the adequate size it is one of the major challenges. The type and concentration of fixative, crosslinking and sonication conditions need to be optimized for each specific tissue. We used a commercially available kit to extract chromatin from cortical microvessels and described the optimized method to isolate and shear chromatin fragments for ChIP assays. In particular, it is critical to properly disrupt the tissue to extract chromatin completely, which generally can be done by pulverizing frozen tissue in liquid nitrogen using mortar and pestle. Due to the challenges of pulverizing a small pellet of microvessels, we first incubated the microvessel preparation in lysis buffer alone, as recommended by the manufacturer’s instructions. However, this procedure was insufficient to completely disrupt them, as it was assessed by the presence of visible structure of intact microvessels in the lysate(Figure 7a). In addition, incomplete disruption yielded very low amount of chromatin. However, when we incorporated a mild sonication step after incubation in lysis buffer, the lysate turned cloudy without any visible intact microvessels (Fig. 7b) resulting in a dramatic increase in the yield.

In addition, we have tested various conditions for crosslinking and sonicating the extracted chromatin and we found that crosslinking with 1% methanol-free formaldehyde for 5 mins, and sonication for 4-8 mins gives rise to 200-400 bp chromatin fragments (Fig. 7c-e), which is the optimal shearing size for most applications.

The condition we provide here may need to be further optimized depending on the levels of expression and the location of the specific DNA binding protein to be pulled down and/or the genomic region/s involved ^30^. However, our data indicate that the sheared chromatin obtained with the provided protocol is suitable for ChIP assay and whole genome sequencing.

## ACKNOLEDGMENTS

This work was supported by funds provided by American Heart Association (Grant-in-Aid, 12GRNT12050110), NIH (HL094465) and Leducq Foundation to T.S.

## AUTHOR CONTRIBUTIONS

HS and TS designed the protocol. HS and YL modified and updated the protocol to its current state. AI conducted stroke surgeries. HY conducted the *in vivo* pharmacological treatments. YL optimized and conducted all the molecular assays with cerebral microvesels. YL, HS and TS wrote the manuscript with contributions of all the authors.

## REFERENCES

1 Zlokovic, B. V. The blood-brain barrier in health and chronic neurodegenerative disorders. Neuron 57, 178–201, doi:10.1016/j.neuron.2008.01.003 (2008).

2 Shi, Y. et al. Rapid endothelial cytoskeletal reorganization enables early blood–brain barrier disruption and long-term ischaemic reperfusion brain injury. Nature communications 7, 10523, doi:10.1038/ncomms10523 (2016).

3 Jackman, K. et al. Progranulin deficiency promotes post-ischemic blood-brain barrier disruption. J Neurosci 33, 19579–19589, doi:10.1523/JNEUROSCI.4318-13.2013 (2013).

4 Bell, R. D. et al. Apolipoprotein E controls cerebrovascular integrity via cyclophilin A. Nature 485, 512–516, doi:10.1038/nature11087 (2012).

5 Montagne, A. et al. Blood-brain barrier breakdown in the aging human hippocampus. Neuron 85, 296–302, doi:10.1016/j.neuron.2014.12.032 (2015).

6 Zhang, Y. et al. An RNA-sequencing transcriptome and splicing database of glia, neurons, and vascular cells of the cerebral cortex. J Neurosci 34, 11929–11947, doi:10.1523/JNEUROSCI.1860-14.2014 (2014).

7 Guo, S. et al. The vasculome of the mouse brain. PLoS One 7, e52665, doi:10.1371/journal.pone.0052665 (2012).

8 Daneman, R. et al. The mouse blood-brain barrier transcriptome: a new resource for understanding the development and function of brain endothelial cells. PLoS One 5, e13741, doi:10.1371/journal.pone.0013741 (2010).

9 He, L. et al. Analysis of the brain mural cell transcriptome. Sci Rep 6, 35108, doi:10.1038/srep35108 (2016).

10 Paolinelli, R. et al. Wnt activation of immortalized brain endothelial cells as a tool for generating a standardized model of the blood brain barrier in vitro. PLoS One 8, e70233, doi:10.1371/journal.pone.0070233 (2013).

11 Ruck, T., Bittner, S., Epping, L., Herrmann, A. M. & Meuth, S. G. Isolation of primary murine brain microvascular endothelial cells. Journal of visualized experiments: JoVE, e52204, doi:10.3791/52204 (2014).

12 Aird, W. C. Endothelial cell heterogeneity. Cold Spring Harbor perspectives in medicine 2, a006429, doi:10.1101/cshperspect.a006429 (2012).

13 Regan, E. R. & Aird, W. C. Dynamical systems approach to endothelial heterogeneity. Circ Res 111, 110–130, doi:10.1161/CIRCRESAHA.111.261701 (2012).

14 Wu, Z., Hofman, F. M. & Zlokovic, B. V. A simple method for isolation and characterization of mouse brain microvascular endothelial cells. Journal of neuroscience methods 130, 53–63 (2003).

15 Munikoti, V. V., Hoang-Minh, L. B. & Ormerod, B. K. Enzymatic digestion improves the purity of harvested cerebral microvessels. Journal of neuroscience methods 207, 80–85, doi:10.1016/j.jneumeth.2012.03.011 (2012).

16 Boulay, A. C., Saubamea, B., Decleves, X. & Cohen-Salmon, M. Purification of Mouse Brain Vessels. Journal of visualized experiments: JoVE, e53208, doi:10.3791/53208 (2015).

17 Yousif, S., Marie-Claire, C., Roux, F., Scherrmann, J. M. & Decleves, X. Expression of drug transporters at the blood-brain barrier using an optimized isolated rat brain microvessel strategy. Brain Res 1134, 1–11, doi:10.1016/j.brainres.2006.11.089 (2007).

18 Kim, G. S. et al. Critical role of sphingosine-1-phosphate receptor-2 in the disruption of cerebrovascular integrity in experimental stroke. Nature communications 6, 7893, doi:10.1038/ncomms8893 (2015).

19 Yanagida, K. et al. Size-selective opening of the blood-brain barrier by targeting endothelial sphingosine 1-phosphate receptor 1. Proc Natl Acad Sci U S A 114, 4531–4536, doi:10.1073/pnas.1618659114 (2017).

20 Kono, M. et al. The sphingosine-1-phosphate receptors S1P1, S1P2, and S1P3 function coordinately during embryonic angiogenesis. J Biol Chem 279, 29367-29373 (2004).

21 Sanchez, T. Sphingosine-1-Phosphate Signaling in Endothelial Disorders. Current atherosclerosis reports 18, 31, doi:10.1007/s11883-016-0586-1 (2016).

22 Zhang, G. et al. Critical role of sphingosine-1-phosphate receptor 2 (S1PR2) in acute vascular inflammation. Blood 122, 443–455, doi:10.1182/blood-2012-11-467191 (2013).

23 Venkataraman, K. et al. Vascular endothelium as a contributor of plasma sphingosine 1-phosphate. Circ Res 102, 669–676 (2008).

24 Sanchez, T. et al. PTEN as an effector in the signaling of antimigratory G protein-coupled receptor. Proc Natl Acad Sci U S A 102, 4312–4317 (2005).

25 Li, Z. et al. Regulation of PTEN by Rho small GTPases. Nat Cell Biol 7, 399–404 (2005).

26 Wolfrum, S. et al. Inhibition of Rho-kinase leads to rapid activation of phosphatidylinositol 3-kinase/protein kinase Akt and cardiovascular protection. Artenoscler Thromb Vasc Biol 24, 1842–1847 (2004).

27 Sanchez, T. et al. Induction of vascular permeability by the sphingosine-1-phosphate receptor-2 (S1P2R) and its downstream effectors ROCK and PTEN. Artenoscler Thromb Vasc Biol 27, 1312–1318 (2007).

28 Ma, D. K. et al. Neuronal activity-induced Gadd45b promotes epigenetic DNA demethylation and adult neurogenesis. Science 323, 1074–1077, doi:10.1126/science.1166859 (2009).

29 Lister, R. et al. Global epigenomic reconfiguration during mammalian brain development. Science 341, 1237905, doi:10.1126/science.1237905 (2013).

30 Nelson, J. D., Denisenko, O. & Bomsztyk, K. Protocol for the fast chromatin immunoprecipitation (ChIP) method. Nature protocols 1, 179–185, doi:10.1038/nprot.2006.27 (2006).

